# The Landscape of Type VI Secretion across Human Gut Microbiomes Reveals its Role in Community Composition

**DOI:** 10.1101/134874

**Authors:** Adrian J. Verster, Benjamin D. Ross, Matthew C. Radey, Yiqiao Bao, Andrew L. Goodman, Joseph D. Mougous, Elhanan Borenstein

**Affiliations:** Departmentof Genome Sciences, University of Washington, Seattle, WA, 98195, USA; Department of Microbiology, School of Medicine, University of Washington, Seattle, WA 98195, USA; Department of Microbial Pathogenesis, Yale University School of Medicine, New Haven, CT 06510, USA; Microbial Sciences Institute, Yale University School of Medicine, West Haven, CT 06516, USA; Howard Hughes Medical Institute, School of Medicine, University of Washington, Seattle, WA 98195, USA; Department of Computational Science and Engineering, University of Washington, Seattle, WA 98195, USA; Santa Fe Institute, Santa Fe, NM 87501 USA

**Author notes:** Equal contribution (Adrian J. Verster, Benjamin D. Ross).

## Abstract

While the composition of the human gut microbiome has been well defined, the forces governing its assembly are poorly understood. Recently, prominent members of this community from the order *Bacteroidales* were shown to possess the type VI secretion system (T6SS), which mediates contact-dependent antagonism between Gram-negative bacteria. However, the distribution of the T6SS in human gut microbiomes and its role have not yet been characterized. To address this challenge, we construct an extensive catalog of T6SS effector/immunity (E-I) genes from three genetic architectures (GA1-3) found in *Bacteroidales* genomes. We then use metagenomic analysis to assess the abundances of these genes across a large set of gut microbiome samples. We find that despite E–I diversity across reference strains, each individual microbiome harbors a limited set of E-I genes representing a single E–I genotype. Importantly, for GA1-2, these genotypes are not associated with a specific species, suggesting selection for compatibility. GA3, in contrast, is restricted to *B. fragilis*, and its low diversity reflects a single *B. fragilis* strain per sample. We further show that in infant microbiomes GA3 is enriched and *B. fragilis* strains are replaced over time, suggesting competition for dominance in developing microbiomes. Finally, we find a strong association between the presence of GA3 and increased abundance of *Bacteroides,* indicating that this system confers a selective advantage *in vivo* in *Bacteroides* rich ecosystems. Combined, our findings provide the first comprehensive characterization of the T6SS landscape in the human microbiome, implicating it in both intra- and inter-species interactions.

## Introduction

Bacterial communities are of fundamental importance to natural ecosystems (Prosser et al. 2007). While cooperative interactions between the species comprising such communities are common (Morris et al. 2012), it is clear that bacteria in these settings also experience pervasive antagonism from surrounding cells (Hibbing et al. 2010). Indeed, the genomes of bacteria encode a wealth of dedicated interbacterial antagonism pathways (Zhang et al. 2012). Some of these function through the production of diffusible small molecules (Riley and Wertz 2002), whereas others utilize proteinaceous toxins. A prevalent pathway mediating the transfer of toxic proteins between bacteria is the type VI secretion system (T6SS). This system has been most thoroughly studied in *Proteobacteria*, though it is found in several phyla of Gram-negative bacteria (Russell et al. 2014a).

The T6S apparatus transfers toxic effector proteins from donor to recipient cells by a mechanism dependent upon cell contact (Russell et al. 2014a). Characterized effector proteins are thus far without exception enzymes that target conserved, essential features of the bacterial cell, such as peptidoglycan, phospholipids, and nucleic acids. This feature of effector proteins, taken together with the fact that T6SS targeting is not dependent on a specific receptor, confers broad activity against Gram-negative cells. Indiscriminate effector transfer also extends to kin cells; therefore, cells with the T6SS produce immunity proteins that inactivate cognate toxins through active site occlusion (Whitney et al. 2015).

Given its wide phylogenetic distribution and its capacity to target diverse recipient cells, the T6SS is likely to play an important role in the assembly and composition of bacterial communities. Indeed, there are recent reports consistent with the pathway mediating bacterial interaction in environmental communities. For instance, T6S genes were found to be enriched and under positive selection in the barley rhizosphere (Bulgarelli et al. 2015), and T6S phospholipase effectors were detected in metagenomes from diverse sources (Egan et al. 2015). To date, however, systematic studies of the impact of T6SS on microbial community assembly are lacking.

The human gut microbiome is a dense ecosystem whose composition is paramount to its function (Walter and Ley 2011). Factors such as diet, immune status, and host genetics have each been implicated in shaping the gut community (Kau et al. 2011), yet the contribution of direct interbacterial competition to the structure of this community remains poorly understood. Recently, a T6SS-like pathway was detected in *Bacteriodetes*, the most abundant Gram-negative phylum in the human gut (Coyne et al. 2014; Russell et al. 2014b). Additional work demonstrated that T6S contributes to the fitness of *Bacteroides fragilis* in competition with other bacteria *in vitro* and in gnotobiotic mice (Russell et al. 2014b; Chatzidaki-Livanis et al. 2016; Hecht et al. 2016; Wexler et al. 2016). These and other data show that the mammalian GI tract is physically conducive to T6S-dependent interbacterial antagonism, suggesting a potential impact of this pathway on the composition of the human gut microbiome (Dong et al. 2013; Sana et al. 2016). Here, we sought to define the distribution of *Bacteroidales* T6S and to explore its function in the human gut microbiome through the analysis of several publicly available metagenomic datasets. These datasets allow us to study the outcome of natural community dynamics in the gut microbiome, and we reasoned that their analysis could therefore provide unique insight into the physiologic role of T6SS-dependent competition in this ecosystem. Our findings reveal the prevalence of this pathway in intact human gut microbial communities, highlight striking and non-random patterns in its distribution across samples, and suggest an active role for T6S in intra- and inter-species interactions in the gut.

## Results

### Detection of T6S E–I pairs in the human gut microbiome

We first set out to characterize the prevalence and distribution of T6SS genes in the gut microbiomes of healthy adult individuals. Based on their organization and content, *Bacteroidales* T6S gene clusters can be divided into three distinct subtypes, termed genetic architecture 1-3 (GA1-3) (Coyne et al. 2016). Each T6S subtype possesses one or more cassettes at stereotyped positions that contain variable genes predicted or demonstrated to encode effector–immunity (E–I) pairs (Russell et al. 2014b; Chatzidaki-Livanis et al. 2016; Coyne et al. 2016; Wexler et al. 2016). As T6S-based antagonism is determined by the effector and immunity genes of donor and recipient cells, respectively, the identification of E–I pairs provides information regarding the potential for interbacterial interactions mediated by this system (Hood et al. 2010). Furthermore, since these cassettes are variable within, but unique among the T6S subtypes, estimation of the abundance of these genes within metagenomes can serve as a proxy for the presence and distribution of GA1-3.

To define the E–I repertoire associated with GA1-3, we searched genes within T6-associated variable cassettes from *Bacteroidales* genomes for those with hallmarks of known T6 effector and immunity factors. These included fusion to modular adaptor domains, reduced GC content, bicistronic arrangement, and similarity to protein families defined by their association with characterized E–I pairs (see Methods for a complete description of annotation criteria; Figures 1A and S1). In total, we identified 9 GA1, 18 GA2, and 14 GA3 putative E–I pairs. As expected, GA3 pairs were identified only in *B. fragilis*, whereas GA1 and GA2 pairs were detected throughout the order. Importantly, we did not identify homologs of GA1-3 E–I genes outside of *Bacteroidales*.

**Figure 1.**
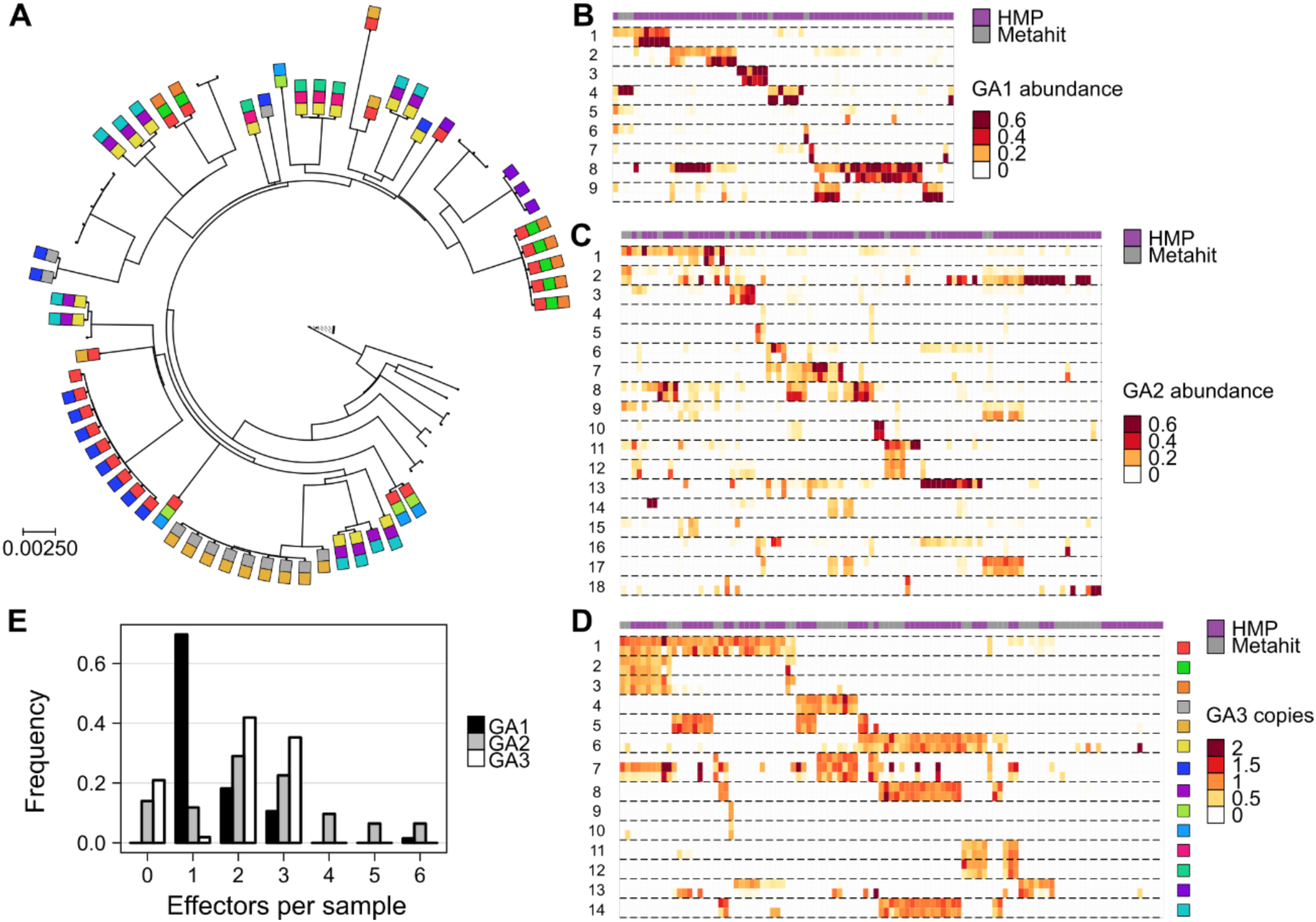
*Bacteroides* T6SS E–I genes are abundant in human gut microbiome samples. **A.** A maximum likelihood phylogeny of *B. fragilis* reference strains constructed from concatenated marker genes. Phylogenetic distance is measured as substitutions per site on the marker genes. GA3 effector genes are represented as colored squares (using the same color coding as in panel D). **B-D.** Each heatmap illustrates the abundance of E–I genes for one of the T6SS subsystems. Each row corresponds to a different E–I pair (effector, top; immunity, bottom). Columns represent the samples analyzed (HMP, purple; MetaHIT, gray). For GA1 and GA2, only samples in which at least 100 reads mapped to the E–I genes of a given subsystem are included, and abundance is measured as fraction of the total abundance of E–I genes in a given sample. For GA3, only samples in which *B. fragilis* is present are included and E–I abundance is normalized by the abundance of *B. fragilis*-specific marker genes, hence measuring the average number of copies per *B. fragilis* genome. **E.** Histograms showing the number of effector genes detected (at >10% of the most abundant effector gene) in each sample.

To estimate the abundance of these E–I pairs in gut microbiomes, we obtained metagenomic datasets derived from healthy donor samples of the HMP (Human Microbiome Project 2012) and MetaHIT (Qin et al. 2010) studies. We next mapped the reads from each sample to the E–I genes using a sequence identity threshold demonstrated to maintain E–I compatibility (Unterweger et al. 2014). Our results indicate that T6S is prevalent in the human gut microbiome; of the 246 samples analyzed, we detected E–I genes in 166 (67%). Moreover, each E–I pair in our list was detected in at least one microbiome sample, with an average of 12.2 occurrences.

### T6S E–I pairs display low diversity within human gut microbiome samples

The systematic characterization of E–I abundance in metagenomic samples provided a unique opportunity to examine the distribution of the genes associated with each genetic architecture across healthy gut microbiomes. We first focused on GA1 and GA2, which utilize unique complements of effectors, but share the ability to undergo conjugative transfer between species belonging to the order *Bacteroidales*(Coyne et al. 2016). Surprisingly, we found that the complement of GA1- and GA2-associated E–I genes in a typical microbiome is small, with only few pairs per sample, comparable to the number of pairs usually detected in a single genome (Figure 1B,C, E). Moreover, in many cases, the same complement of E–I genes was detected in multiple samples. Henceforth, we refer to these combinations as E–I genotypes. This pattern suggests that either each sample is dominated by a single strain that harbors the observed E–I genotype or that there exists selective pressure for compatible E–I genes across multiple strains or species in a sample.

To further explore these possibilities, we focused our attention on the most prominent members of the genus *Bacteroides*. Other genera in the order *Bacteroidales* are less abundant constituents of the microbiome and based on reference genomes do not often harbor GA1 or GA2. We identified a set of species-specific single-copy marker genes for each *Bacteroides* species and estimated their abundance in each sample. Next we compared marker gene abundance to that of GA1 and GA2 E–I genes across samples (see Methods). We found that the abundance of these E–I genes was not consistent with that of any individual species (Figures 2A and S2). These findings suggest that multiple species co-existing in a microbiome typically encode a single GA1 and/or GA2 E–I genotype, potentially due to selective pressure for maintenance of E–I compatibility.

**Figure 2.**
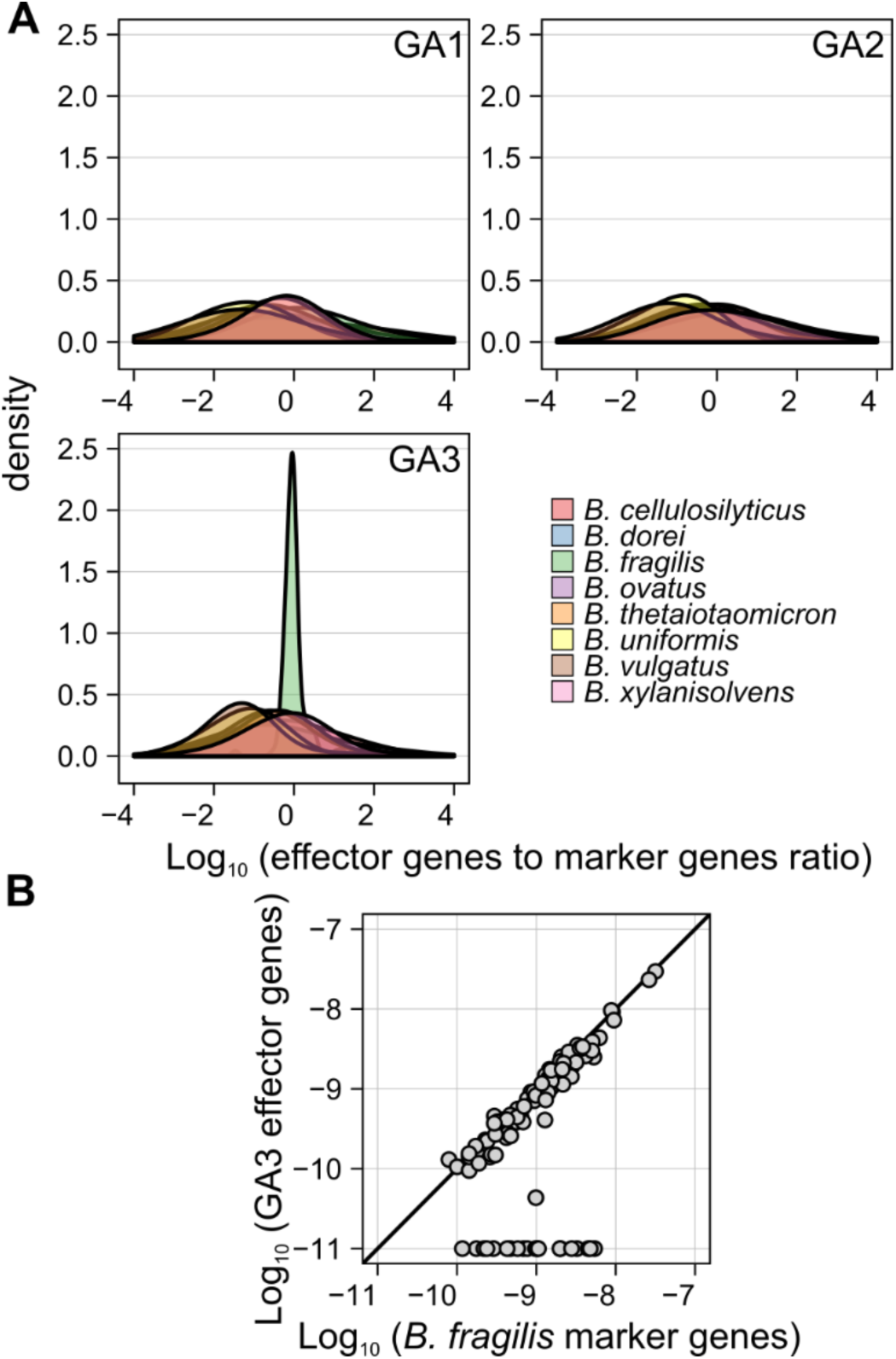
Differential associations between T6SS and *Bacteroides* spp. **A.** Density plots showing the distribution across samples of the ratio between the average abundance of detected effector genes from each T6SS subsystem and the average abundance of species-specific marker genes for different *Bacteroides* spp. Only samples in which at least 100 reads mapped to the E–I genes of a given subsystem and only species for which at least 5 genomes were available (and therefore marker genes can be robustly inferred) are included. **B.** Scatter plot of the average abundance of detected GA3 effector genes vs. the average abundance of *B. fragilis*-specific marker genes. Only samples in which *B. fragilis* is present are included. A small factor (10^−11^) has been added to each abundance value to allow transformation to a logarithmic scale.

We next examined GA3 E–I genes, and found, as in GA1 and GA2, that each sample harbors only a small set of E–I pairs (Figure 1D,E). Moreover, observed GA3 E–I genotypes matched those detected in reference genomes (Figure S3), and appeared randomly distributed between the American (HMP) and European (MetaHIT) datasets (Figure 1D). However, in contrast to GA1 and GA2, we found a strong correlation (R = 0.94) between the abundance of GA3 effector and immunity genes and that of a single species, *B. fragilis* (Figures 2A-B and S2). This finding is supported by our curation of E–I genes and by previous studies (Coyne et al. 2016) that found GA3 restricted to *B. fragilis*. Importantly, however, our finding confirms that this restriction of GA3 to *B. fragilis* observed in sequenced reference genomes holds across naturally occurring communities.

We hypothesized that the pattern of GA3 E–I genotypes we observed could be explained by the dominance of a single *B. fragilis* strain within each individual microbiome. Indeed, prior studies suggest that *B. fragilis* exhibits relatively low diversity within individuals (Yassour et al. 2016). To confirm that this pattern is also observed in HMP and MetaHIT samples, we first measured nucleotide diversity in species-specific markers of *Bacteroides* spp. We found that, within an individual, *B. fragilis* possesses the lowest average SNP diversity of well represented members of the *Bacteroides* genus (Figure S4A). We then used a previously developed method for inferring the most likely set of strains in metagenomic samples based on nucleotide variants, combined with a phylogenetic analysis of these inferred strains to determine the number of different monophyletic groups of strains present in each sample (see Methods). We found that *B. fragilis* inferred strains in every HMP and MetaHIT sample formed a single monophyletic group, indicating that extant *B. fragilis* strains in each sample are likely derived from a single colonization event. Moreover, the set of E–I pairs detected in each metagenomic sample generally matched the set of E–I pairs found in the reference strains closest in the phylogenetic tree to the inferred strains, especially when the distance of inferred strains to their nearest reference was low (Figure S5).

To experimentally confirm our computational findings, we additionally selected 20 randomized *B. fragilis* colonies isolated from two healthy adults and subjected these to whole genome sequencing. Consistent with our findings using metagenomic data, our sequencing showed that a single clonal strain of *B. fragilis* dominates the microbiome of these individuals (Figure S4B).

### *B. fragilis* GA3 is important in the developing microbiome

The finding that the presence of singular GA3 genotypes within individuals is due to the dominance of one *B. fragilis* strain motivated us to investigate the role of this system in the microbiome. We reasoned that in this dense and competitive microbiome ecosystem (Ley et al. 2006), an antagonistic pathway such as the T6SS might provide a fitness advantage. The system could mediate antagonism against other *B. fragilis* strains closely related organisms such as other *Bacteroides* spp., other Gram-negative inhabitants of the microbiome, or a combination of these (Chatzidaki-Livanis et al. 2016; Hecht et al. 2016; Wexler et al. 2016). Assuming such a role for T6SS, we further reasoned that in the microbiome of infants, which is less stable than that of adults(Sharon et al. 2013), the function of an antagonistic pathway like the T6SS might be more pronounced. To test this hypothesis, we obtained publically available metagenomic datasets derived from infant gut microbiomes (Backhed et al. 2015; Kostic et al. 2015; Vatanen et al. 2016; Yassour et al. 2016). We then identified samples that contain *B. fragilis* but lack GA3-associated structural genes (see Methods) in both adult and infant datasets. Such samples indicate the presence of *B. fragilis* strains unable to intoxicate competitor bacteria using this pathway. The availability of well-assembled *B. fragilis* reference genomes that lack the GA3 gene cluster lent credence to this approach (Wexler et al. 2016). We found that infant microbiomes containing *B. fragilis* are significantly less likely to lack GA3-associated structural genes relative to those of adults (Fisher’s exact test, P < 0.01, 8% infants, 23% adults; n = 276; Figures 3A and S6).

**Figure 3.**
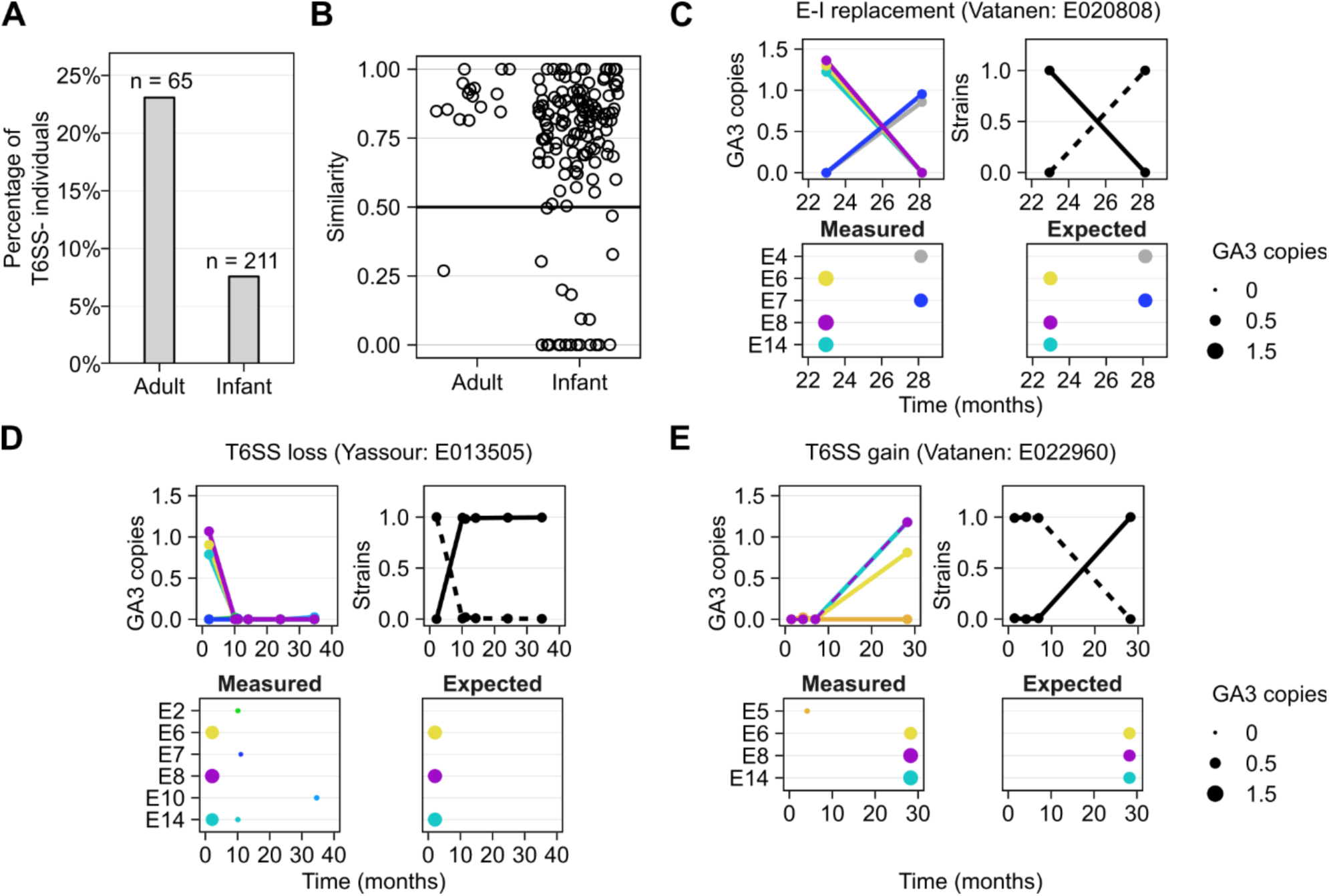
E–I turnover and strain replacement in infant microbiomes. **A.** The percentage of individuals of those harboring *B. fragilis,* that lack the GA3 T6SS across adult and infant datasets. **B.** The minimal similarity (measured by the Jaccard similarity coefficient) in GA3 E–I gene content between the first time point and every subsequent time point in adults and infants. **C-E.** Examples of E–I turnover events and corresponding strain replacement events are shown. The plots on the upper and bottom left in each panel illustrate the estimated abundance of GA3 effector genes (measured as copies per *B. fragilis* genome) over time, with the plot on the upper right illustrating the estimated frequency of inferred strains in these samples. Only samples in which *B. fragilis* is present are shown. The bottom right plot illustrates the expected abundance of the various effector genes based on the effector genes encoded by reference strains that are phylogenetically close to the inferred strains.

This finding suggests that GA3 provides an advantage for *B. fragilis* in early life; however, the selection pressure underlying this advantage remained unclear. Several independent studies using gnotobiotic mice have shown that the GA3 T6SS can play a major role in the competition between *B. fragilis* strains in the gut (Chatzidaki-Livanis et al. 2016; Hecht et al. 2016; Wexler et al. 2016). However, *B. fragilis* is thought to be stable after acquisition from the mother, and inter-strain competition within the human gut microbiome has not been documented for this organism (Faith et al. 2013; Nayfach et al. 2016). Aiming to capture such processes in the developing microbiome, we estimated the abundance of GA3 E–I genes for individual infant samples as we did for adults. In general, the E–I landscape of infants mirrors that of adults, with generally a single genotype present in each sample (Figure S7). Moreover, many of the most prevalent E–I genotypes we observed in adults are also frequent in infants.

Notably, the infant microbiome datasets we analyzed include multiple samples per individual, thereby allowing us to examine the temporal dynamics of *B. fragilis* and of T6SS genes. Surprisingly, this analysis revealed many instances in which the E–I genotype of an individual has changed between samples (Figure 3B). In total, we observed E–I turnover in 22 of the 117 infants for which longitudinal data was available. Such E–I turnover events include instances where one GA3 genotype is replaced by another (Figure 3C), but also gains and losses of the T6SS (Figure 3D-E). To further confirm these E–I dynamics, we used the strain inference method described above. We detected a corresponding strain replacement in 17 of the 22 individuals in which an E–I turnover event was observed (Figures 3C-E and S8). Moreover, comparing the set of E–I genes detected in each sample to those encoded by the reference strains phylogenetically closest to the inferred strain, we further find overall agreement between observed and expected E–I turnover events. Notably, instances of one strain replaced by another with a similar E–I genotype (Figure S8; Vatanen:T014827) or of transient co-existence of E–I genotypes (Figure S8; Backhed:587) were also observed. Interestingly, examining the few HMP adult individuals for which data was available from multiple visits, we found one adult in which the E–I genotype similarly changed over time (Figure 3B).

### *B. fragilis* GA3 T6SS is associated with shifts in community composition

Due to its lower frequency in adult microbiomes compared to those of infants, the GA3 T6SS is absent in many adult samples in which *B. fragilis* can be detected (23%), offering a unique opportunity to compare the community composition in samples with or without GA3, and to uncover community-associated outcomes of T6S activity in the human gut. To this end, we obtained the taxonomic profile of all HMP samples (see Methods) and identified associations between these profiles and the presence of T6SS structural genes. We first compared overall community composition between samples as measured by the Bray-Curtis distance. We found that samples harboring *B. fragilis* and GA3 genes (T6SS+) significantly differ in community composition from samples harboring *B. fragilis* but lacking these genes (T6SS-; P < 0.01 PERMANOVA; n = 51). Examining the abundance of each genera across samples, we further identified four genera whose abundance in T6SS+ versus T6SS-samples significantly differs (Wilcoxon rank sum test; FDR<0.05; Figure 4 and Table S1). Specifically, we found that the abundance of *Bacteroides* is positively correlated with the presence of GA3, which is consistent with experimental and theoretical work indicating that members of this genus are most likely to compete with *B. fragilis* for its niche (Russell et al. 2014b; Trosvik and de Muinck 2015; Chatzidaki-Livanis et al. 2016; Hecht et al. 2016; Wexler et al. 2016). Furthermore, the genera *Faecalibacterium*, *Oscillospira* and *Ruminococcus* from the phylum *Firmicutes* were negatively correlated with GA3. Gram-positive organisms are not targets of any known T6SS; therefore, the observed decreases in abundance of these genera in T6SS+ microbiomes are likely to be the indirect result of selection for GA3 occurring in communities with an increased ratio of *Bacteroidetes* to *Firmicutes*.

**Figure 4.**
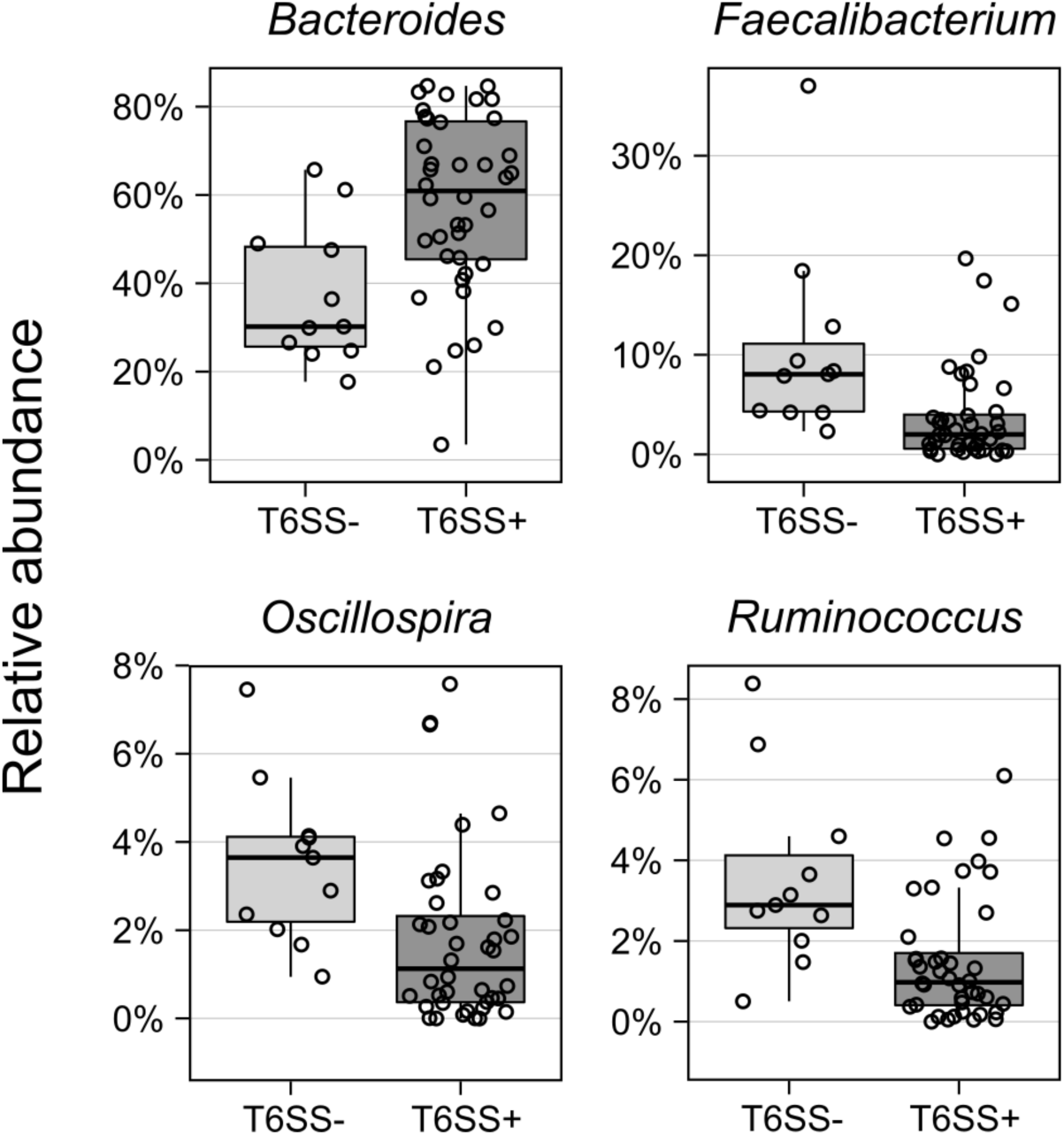
Differentially abundant genera between T6SS+ and T6SS- HMP samples. Abundances are based on a 16S rRNA survey and only genera whose abundances are significantly different in T6SS+ vs. T6SS- samples (at FDR < 0.05) are plotted.

## Discussion

Despite the wide distribution of T6S in Gram-negative bacteria, little is known about its role in natural communities. Here, we surveyed human microbiome samples and discovered that these communities are replete with bacteria containing the T6SS. We focused our analyses on genes specific to the order *Bacteroidales*; thus, our findings are an underestimate of the prevalence and impact of this system in gut communities. Nonetheless, our characterization of T6SS E–I gene distribution in human gut microbiomes suggests that this contact-dependent pathway plays a role in competition and selection at multiple levels.

We observed a markedly low diversity of T6SS E–I genes in human microbiome samples. Specifically, a single genotype of GA1 and GA2 E–I genes is found in each microbiome, yet the abundance of these genes does not correlate with that of any one species in the *Bacteroides* genus. This supports a model in which antagonism via GA1 and GA2 exerts selective pressure for compatibility between *Bacteroides* spp. in the gut. We postulate that in the case of GA1 and GA2, horizontal transfer facilitates E–I compatibility. Indeed, Comstock and colleagues found that transfer of GA1 and GA2 can occur between species within the microbiome of an individual (Coyne et al. 2014).

Similar to GA1 and GA2, we found a single genotype of GA3 in each microbiome, but were able to demonstrate that GA3 is restricted to *B. fragilis* and can be explained by the presence of a single *B. fragilis* strain in each individual. This is consistent with experimental studies demonstrating a role for GA3-dependent competition between *B. fragilis* strains (Chatzidaki-Livanis et al. 2016; Hecht et al. 2016; Wexler et al. 2016). We found, however, that infant microbiomes commonly exhibit a strain replacement accompanied by E–I turnover.

While the low number of adult longitudinal samples prohibits us from statistically comparing strain replacement rates in infants and adults, the observed strain replacements in infants and our finding that *B. fragilis* lacking the GA3 T6SS is more common in adults suggest that, early in life, *B. fragilis* strains compete for dominance, potentially contributing to the establishment of this system in the infant microbiome. This is further supported by the fact that we rarely find co-existing strains of *B. fragilis,* and that typical turnover events are characterized by the rapid transition of one dominant strain to another. Indeed, gnotobiotic experiments indicate that inter-strain competition is strong in mice colonized solely with *B. fragilis,* and diminished when the relative abundance of *B. fragilis* is reduced in a more complex community (Wexler et al. 2016). Therefore, the infant microbiome may represent a particularly dynamic ecosystem in which the GA3 T6SS facilitates *B. fragilis* strain competition.

We find that strains of *B. fragilis* lacking GA3 are more commonly found in adults than infants. This could arise either by the replacement of T6SS+ with T6SS- strains, or by the loss of the T6SS system from a previously T6SS+ strain of *B. fragilis*. This decline in *B. fragilis* GA3 prevalence in adulthood may reflect a change in its selective advantage. Indeed, there is precedent for the lability of T6S in bacteria undergoing strong shifts in environmental context, such as *Burkholderia mallei* and *Bordetella* spp. (Schwarz et al. 2010). It is likely that community effects buffering the *B. fragilis* niche develop with the maturation of the more stable adult gut community. Stabilization over development appears to render GA3 dispensable in certain contexts, for instance within those microbiomes that contain lower populations of potential *B. fragilis* competitors and known targets of the GA3 pathway, other *Bacteroides* spp. Our findings and others suggest that *B. fragilis* primarily faces selective pressure from closely-related species (Chatzidaki-Livanis et al. 2014; Russell et al. 2014b; Chatzidaki-Livanis et al. 2016; Hecht et al. 2016; Roelofs et al. 2016; Wexler et al. 2016). Put differently, the observed association between GA3 and community assembly reflects selection on GA3 mediated by community composition, rather than GA3 mediated impact on overall community assembly. Importantly, *B. fragilis* abundance is low compared to the *Bacteroides* consortium as a whole and therefore, it is perhaps not surprising that we do not detect other Gram-negative genera whose abundance are specifically lowered in GA3+ microbiomes. However, we cannot rule out that competition from less common or low abundance genera that fall below our detection limit might also select for the retention of GA3 in adults. Indeed, there is evidence that *Bacteroides* spp and members of the *Enterobacteriaceae* interact intimately, although T6SS- dependent interactions have not yet been shown between these organisms (de Sablet et al. 2009; Buffie and Pamer 2013; Curtis et al. 2014; Charbonneau et al. 2016).

The T6SS is one of many antagonistic pathways whose operation is determined by the presence or absence of polymorphic toxins and corresponding antitoxins (Aoki et al. 2010; Zhang et al. 2012; Cao et al. 2016). We show here that a systematic characterization and large-scale computational analysis of metagenomic data can provide a means of linking the presence and abundance of these crucial factors to microbial community composition. Moreover, for contact-dependent pathways such as the T6SS, such analyses can provide a unique window into community biogeography. Indeed, *Bacteroides* spp. are thought to occupy a crowded niche proximal to the gut mucosa and our findings herein provide evidence of extensive cell–cell contacts between species of the genus (Earle et al. 2015; Donaldson et al. 2016). Thus, our study offers an analytical framework for more globally deciphering the forces that dictate the establishment and maintenance of bacterial communities.

## Methods

### Short read metagenomic datasets and genomic data

Our analysis utilizes short read metagenomic data from several large-scale microbiome datasets. For adult microbiomes we downloaded 147 shotgun samples from HMP (Human Microbiome Project 2012), and 99 healthy human shotgun samples from MetaHIT (Qin et al. 2010). Since an excessive fraction of human DNA will likely not markedly impact our ability to quantify *B. fragilis* abundance, HMP samples which failed QC were nonetheless included in our analysis. For infant microbiomes, we downloaded 300 samples from a study of development of the microbiome in the first year of life (Backhed et al. 2015), 769 samples from a study of autoimmune diseases (Vatanen et al. 2016), 237 samples from a study of antibiotic usage (Yassour et al. 2016), and 126 samples from a study of the development of Type 1 Diabetes (Kostic et al. 2015). Several of these datasets include multiple longitudinal samples from the same individuals, which were used for temporal analysis.

We downloaded all available *B. fragilis* genomes from RefSeq. Sequences from 3 strains were found to be contaminated with contigs matching species other than *B. fragilis*, and were discarded. A group of 8 strains appeared to be very distant in sequence homology from the rest of the strains, and were also discarded. We additionally downloaded from RefSeq all genomes of other *Bacteroides* species for which at least 5 strain genomes were available.

### Identifying species-specific marker genes

We compile a list of marker genes that could be used for strain-level inference. Our marker gene approach is similar to that used by MetaPhlAn (Truong et al. 2015), but rely on a more stringent selection of marker genes, supporting a more robust comparison at the strain level. Specifically, for our analysis, we identified a subset of the MetaPhlAn marker genes that are found in the genome of *every* sequenced strain in a single copy. To this end, for each of the MetaPhlAn marker genes associated with a given species, we used BLASTn to find every homolog (>60% identity) in all the strains of that species. We used Usearch (Edgar 2010)to cluster this larger set of genes into groups with >90% identity and if a cluster with exactly one gene in each strain could be found, the marker gene was included in our list.

### Identifying T6SS effector and immunity genes

We sought to comprehensively catalog *Bacteroidales* T6SS E–I genes from reference genomes. In *Bacteroidales*, as in other bacteria, E–I genes are encoded adjacent to the genes for secreted structural proteins Hcp and VgrG. Accordingly, we manually curated genes adjacent to these structural genes across all publically available *Bacteroidales* genomes. Identified genes exhibited reduced GC content relative to the rest of the T6SS locus or the genome as a whole, and were encoded in bi-cistrons. Putative effectors always lacked characteristic signal peptides, consistent with transport via the T6SS apparatus, while putative immunity genes often encoded proteins with signal peptides. We used structural homology prediction (Phyre)(Kelley et al. 2015) and remote sequence homology search algorithms (Hmmer)(Finn et al. 2015) to predict functions for these genes, identifying many genes with functions associated with known T6SS toxin effectors. As in Coyne *et al* (Coyne et al. 2016), our list included predicted cell-wall degrading enzymes, lipases, and nucleases as well as putative effector domains fused to either PAAR domains (DUF4280) or Hcp.

### Estimating the abundance of species-specific genes and T6SS genes in metagenomic samples

We aligned shotgun reads single end using Bowtie2 (using parameters –a –N 1) to the set of genes of interest. Alignments with less than 97% identity, a quality score below 20, or multiple hits were discarded. To quantify the abundance of each gene, the number of reads aligned to this gene was normalized by the length of the gene and the total number of reads in the sample.

The average abundance of species-specific marker genes identified above was used as a proxy for the abundance of that species in the sample. We defined samples as having *B. fragilis* present if at least 100 reads could be aligned to *B. fragilis*-specific marker genes. When characterizing strain replacement, for which higher coverage of *B. fragilis* genes is required, we used instead a threshold of 500 reads. Because GA1 and GA2 are not restricted to a single species, and because *Bacteroidales* composition can vary dramatically, we considered samples with 100 reads mapping to the GA1 and GA2 E–I genes.

### Nucleotide diversity calculation

To estimate nucleotide diversity across species-specific marker genes, we again aligned all short reads in each sample to these genes. The obtained alignments were converted into a pileup using mpileup from samtools (parameters --excl-flags UNMAP,QCFAIL,DUP -A -q0 -C0 –B), and finally into an allele count matrix. The first and last 10 bases of each gene were discarded from the allele count matrix as we found they contained many poor quality alignments. We focused on high-coverage loci only, ignoring all loci where the coverage was less than 5X. If the number of high coverage sites was <10% of the total length of the sequence, the sample was excluded from further consideration. Variable sites were defined as those having at least 2 counts of the minor allele. Nucleotide diversity was then calculated at these variant sites according to:

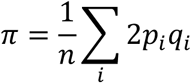

Where *n* is the total length of the genes, *i* corresponds to the variable sites, and *p* and *q* correspond to the frequency of the major and minor allele at site *i*.

### Inferring *B. fragilis* strains using StrainFinder

To infer strain diversity in each sample, we used StrainFinder (https://github.com/cssmillie/fmt), a previously introduced method for inferring the most likely set of strains in metagenomic samples based on nucleotide variants. To this end, we again aligned the short read in each sample to the set of *B. fragilis*-specific marker genes identified above, and converted the alignment to a count matrix describing the number of counts of each nucleotide at every position along the genes. As when calculating nucleotide diversity, we discarded the first and last 10 bases and only considered sites with at least 5 counts. We also discarded samples where the high coverage sites were less than 10% of the total length as they resulted in poor quality trees. When running StrainFinder we reduced the data to only those sites with population variability, defined, as above, as sites with at least 2 counts of the minor allele. As noted above, we only considered samples with a sufficient coverage on the *B. fragilis* marker genes to enable robust strain inference.

StrainFinder determines the relative strain abundance and genotypes at variable sites by considering the likelihood of the observed allele counts and using an expectation maximization approach. An optimal number of strains between 1 and 10 was determined using AIC. For each run of StrainFinder we used 5 independent runs of 200 expectation maximization iterations and selected the best fit; these parameters yield reproducible inference of strains. When analyzing temporal data with StrainFinder, we combined allele counts from all samples of an individual into a single 3-dimensional matrix. Using the genotypes from StrainFinder we then reconstructed strain-specific versions of each marker gene and then created subsequences consisting of only the high coverage sites we used in the analysis. Inferred strains were then further examined using a phylogenetic analysis as described below.

### Phylogenetic analysis

A phylogenetic tree of the reference *B. fragilis* strains was constructed based on their species-specific marker genes. Specifically, we aligned the strains’ versions of each marker gene using MAFFT, concatenated the alignments of all genes, and then constructed a tree using the GTRGAMMAI model from RAxML (Stamatakis 2014), as has been done previously (Wexler et al. 2016).

To determine whether the strains inferred by StrainFinder are monophlyletic, we combined the sequences from the inferred strains (as determined by StrainFinder), with the sequences from the available reference *B. fragilis* genomes, and recreated the strain phylogeny using the same method as described above. We defined inferred strains as monophyletic if their common ancestor does not have any descendent outside the set of inferred strains or if the distance was less than 0.001 substitutions per site.

### Strain sequencing

All human studies were conducted with the permission of the Yale Human Investigation Committee. Stool samples from four healthy individuals frozen in sterile glycerol (Cullen et al. 2015) were plated onto *Bacteroides* Bile Esculin or *Brucella* Blood Agar plates (BD Biosciences), to select for *Bacteroidales* colonies. Single colonies were picked into Mega Medium (Wu et al. 2015) and grown to stationary phase in anaerobic conditions before freezing in 10% glycerol in 96-well plates. PCR was performed directly from the frozen stocks using primers to amplify the V1-V4 region of the 16S rRNA gene. PCR products were then sent for Sanger sequencing. Reads were converted to fastq format and NCBI Blast 2.2.31+ was used to align sequences to the SILVA 123 and GreenGenes 2011-1 16S rRNA gene databases in order to identify *Bacteroides fragilis* colonies. To verify the *B. fragilis*-positive colonies, a second round of PCR was performed using primers to amplify and sequence the gyrB gene. Two of the four donors were confirmed to have *B. fragilis*. Twenty confirmed colonies from each *B. fragilis*-positive donor were then grown up to stationary phase in TYG medium under anaerobic conditions. Genomic DNA was isolated using the Qiagen DNeasy Blood and Tissue Kit and prepared for whole genome sequencing using the MiSeq V3 Reagent Kit. Sequencing was performed in the Nickerson lab core facility in the UW Department of Genome Sciences. Sequencing reads were mapped to the set of *B. fragilis*-specific marker genes to generate alignments. Samples under 10x mean alignment read coverage were then discarded. Consensus sequence for each remaining sample was generated using the GATK FastaAlternateReferenceMaker. Subsequently, we constructed multi-alignments for all the samples using MAFFT 7.237, concatenated them, and then inferred a phylogenetic tree using the GTRGAMMAI model from RAxML 8.2.8 (Stamatakis 2014). All sequences were deposited into NCBI SRA under BioProject ID PRJNA375094.

### Predicting E–I gene content of inferred *B. fragilis* strains

We predicted the E–I gene content of an inferred *B. fragilis* strain by examining the E–I content in the genome of its nearest neighbors on a phylogenetic tree that contains the inferred strains from a given sample and the reference strains (as described above). Specifically, for every strain identified from StrainFinder, we identified the most recent ancestor that have both the inferred strain and at least one reference strain as descendants. We then used the average E–I content of all reference strains descendant from this ancestor as the predicted E–I content of the inferred strain. To then estimate the predicted E–I content in the sample, we combined the predicted E–I content of each inferred strain weighted by their relative abundance. To determine the confidence of the predicted E–I content we determined the average phylogenetic distance of these reference strains to the ancestor identified above.

### Classifying microbiome samples as T6SS+ vs. T6SS-

For every sample, we estimated the number of reads expected to map to the *B. fragilis* GA3 T6SS structural genes based on the number of reads mapped to *B. fragilis*-specific marker genes in that sample and the ratio between the total length of *B. fragilis*-specific marker genes and *B. fragilis* T6SS structural genes. We define samples to be T6SS+ if *B. fragilis* was present (as defined above) and the number of reads mapped to T6SS structural genes was more than 10% of the expected number (and see Figure S6A). We define samples to be T6SS- if *B. fragilis* was present and the number of reads mapped to T6SS structural genes was less than 10% of the expected number.

### Community composition analysis

To obtained independent estimate community composition in each sample, we downloaded the v35 16S OTU abundance table for human gut microbiomes from HMP (ftp://public-ftp.hmpdacc.org/HMQCP/otu_table_psn_v35.txt.gz), summed the counts from all OTUs in the same genus, and calculated the relative abundance of each genus. Importantly, because 16S sequencing depth is independent from the depth of shotgun samples used to determine T6SS+ vs. T6SS- classification, using these 16S-based data allows us to compare T6SS presence with community taxonomic profiles without potential coverage-related biases. Samples were classified into T6SS+ and T6SS- as described above. GA1 and GA2 lack uniquely identifying structural genes so we defined T6SS+ vs. T6SS- as samples with vs. without 100 counts mapping to the GA1 or GA2 E– I genes respectively. The distance between samples was defined by the Bray-Curtis distance at the genus level, and significance of separation between T6SS+ and T6SS- samples was evaluated using PERMANOVA. For the subset of genera whose average abundance across samples was > 0.1%, we used a Wilcoxon rank sum test to compare their abundance in T6SS+ vs. T6SS- samples using a 5% FDR.

## Acknowledgements

We thank Eric Alm and Chris Smillie for sharing StrainFinder code and for their support in running it. We also thank UW Genome Sciences ITS for high-performance computing resources. We are grateful to S. Brook Peterson for careful review of the manuscript, and to members of the Borenstein and Mougous laboratories for helpful discussions. This work was supported by National Institutes of Health grants GM118159 (to ALG), AI080609 (to JDM), and New Innovator Award DP2AT00780201 (to EB), the Pew Scholars Program (ALG), and the Burroughs Wellcome Fund (ALG and JDM). AJV was supported by a postdoctoral fellowship from the Natural Sciences and Engineering Research Council of Canada. BDR was supported by a Simons Foundation-sponsored Life Sciences Research Foundation postdoctoral fellowship.

## Supporting Figure Legends

**Figure S1.**
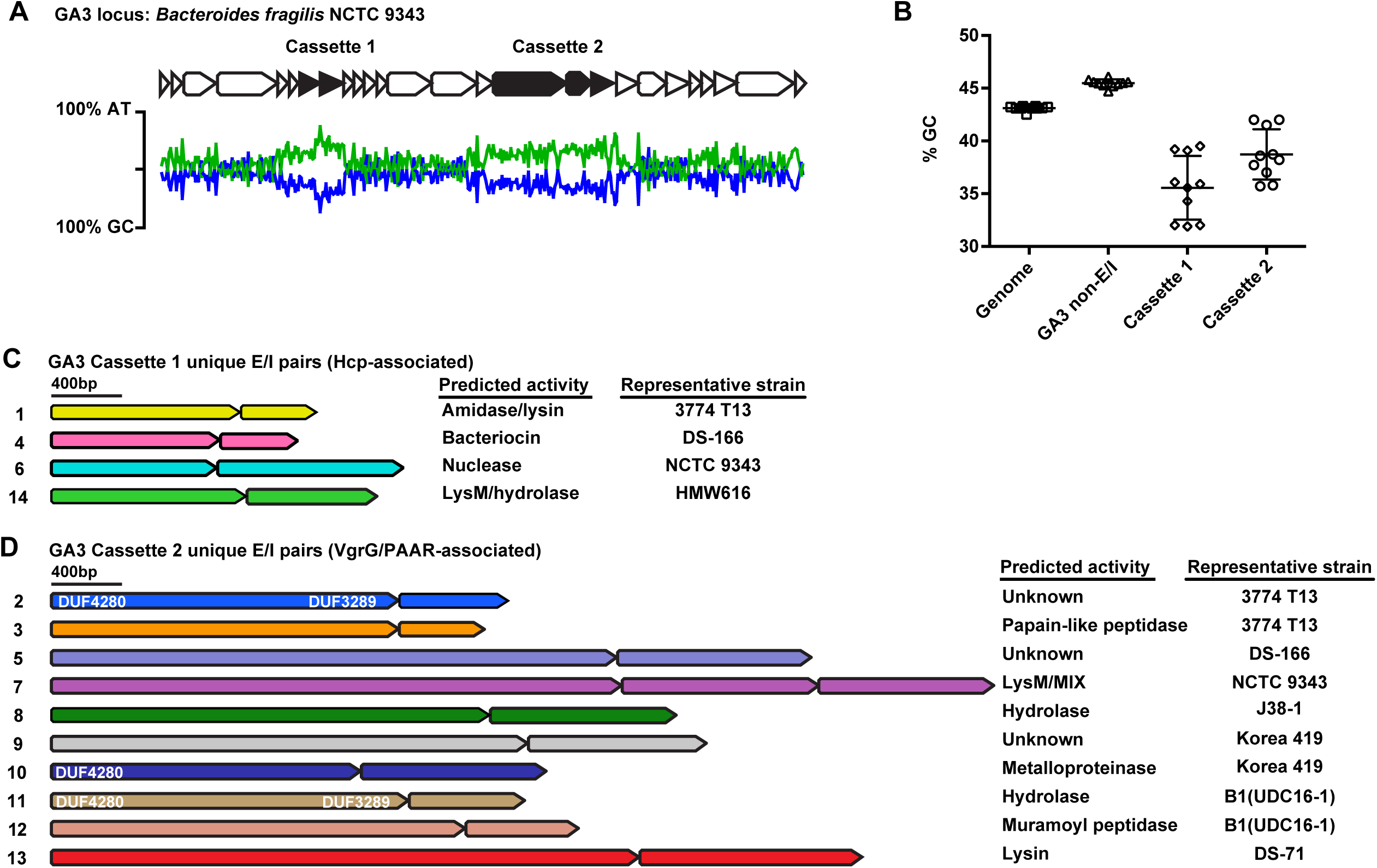
Identification of unique *B. fragilis* GA3 effectors. **A.** Schematic of the GA3 locus from *B. fragilis* NCTC 9343. GC and AT content plotted below locus. **B.** GC content of indicated regions from 10 representative *B. fragilis* strains. **C.** Unique *B. fragilis* GA3 E–I pairs from cassette 1 and cassette 2 used in subsequent metagenomics analyses.

**Figure S2.**
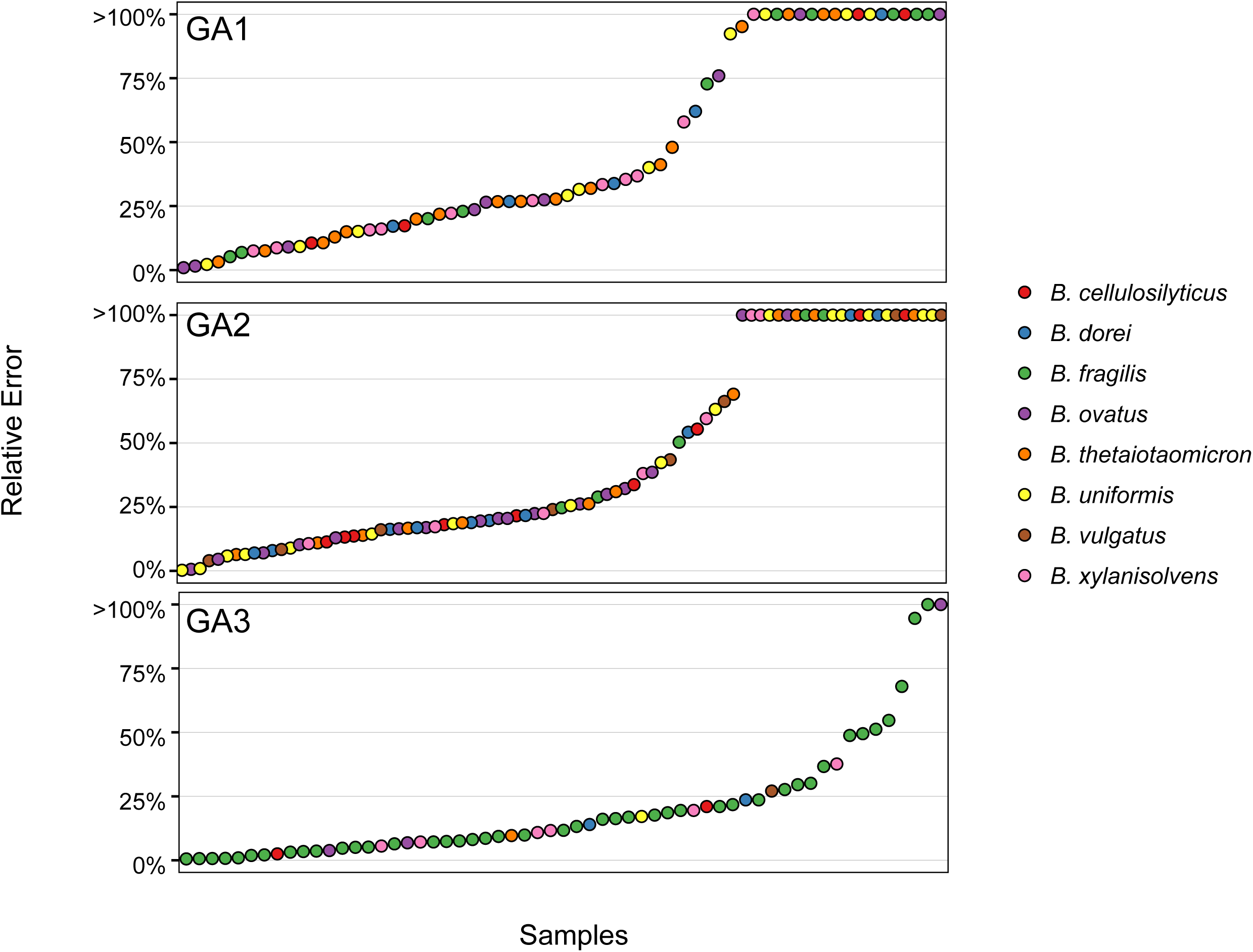
Minimal relative error in effector abundance assuming that the T6SS is encoded by a single species. The relative error is defined as the difference between the average abundance of detectable effector genes in a sample and the expected abundance of these genes assuming they are encoded by a given species. For each sample, the minimal relative error (across all possible species) is plotted and samples are ordered by the magnitude of the minimal relative error. Only samples in which at least 100 reads mapped to the E–I genes of a given subsystem are included. The color of each point represents the species for which the minimal relative error was obtained.

**Figure S3.**
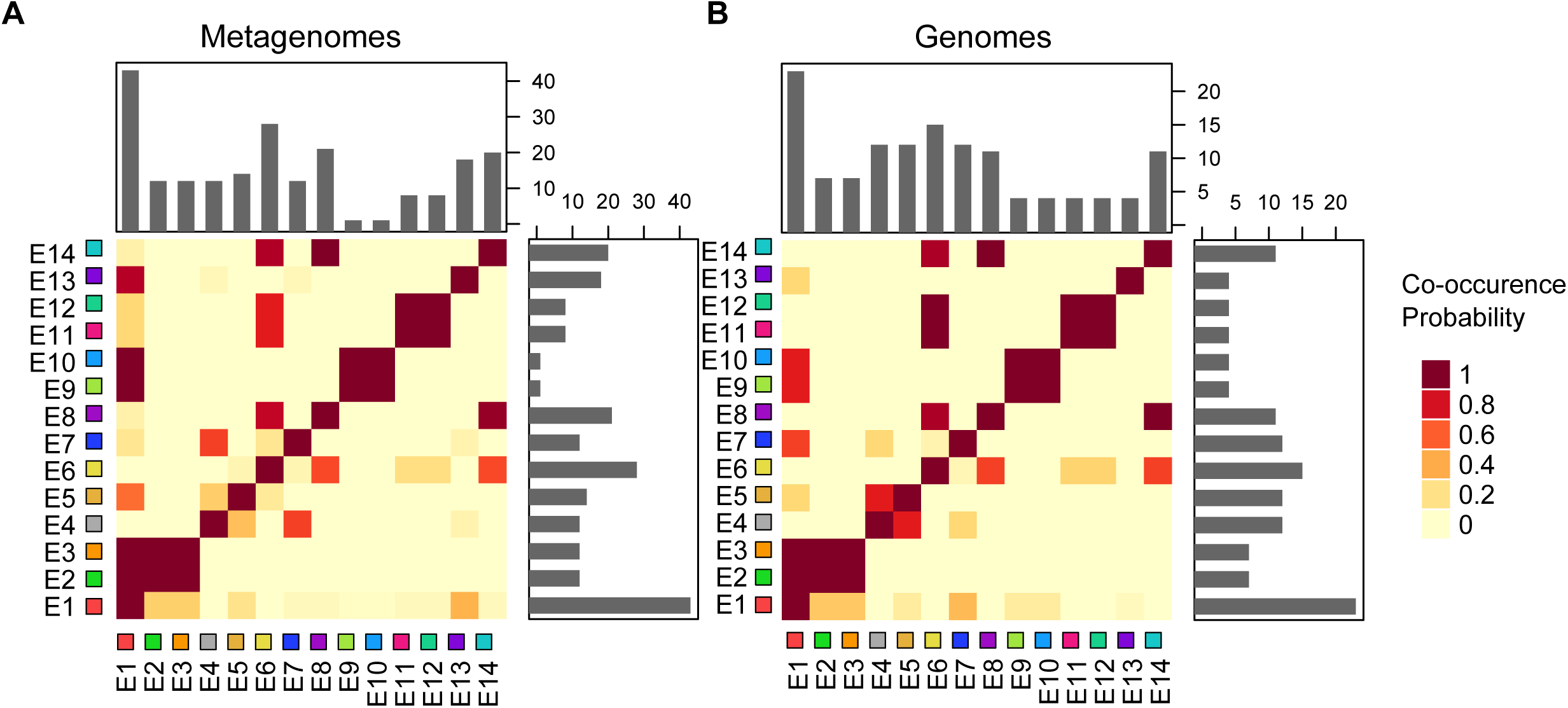
The co-occurrence of GA3 effector genes in metagenomes and genomes. Each cell in the heatmap, *a*_*ij*_, denotes the probability that effector gene *i* is detected in a sample **A.** or encoded in a genome **B.** given that effector gene *j* was detected/encoded. The barplots on the top and right of each heatmap illustrate the number of metagenomes or genomes in which each effector gene was detected.

**Figure S4.**
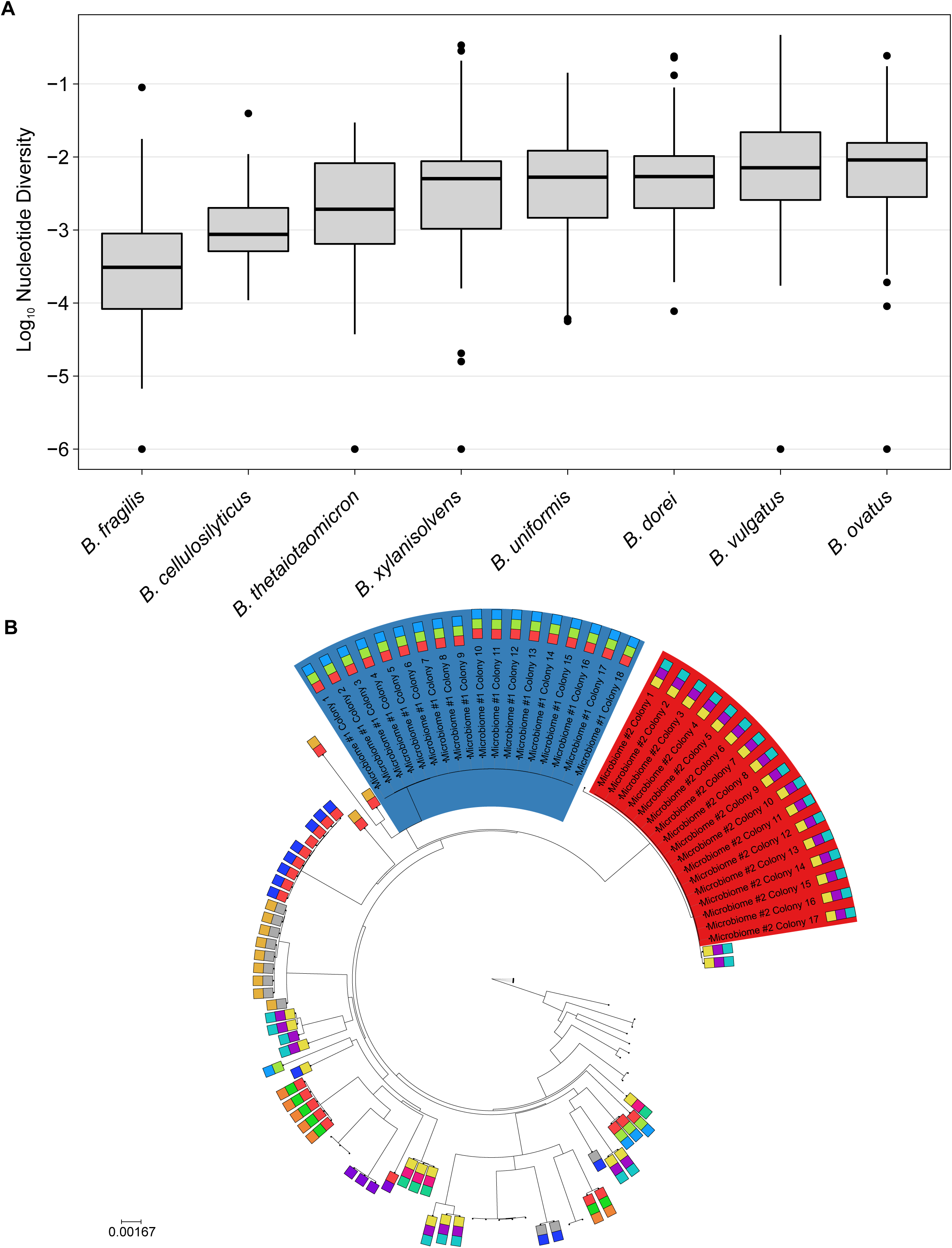
*Bacteroides* nucleotide and strain diversity in the gut microbiome of adults. **A.** The nucleotide diversity for different *Bacteroides* spp. in adult samples from the HMP and MetaHIT studies. Nucleotide diversity was calculated based on population variants in species-specific marker genes (Methods). Only species with at least 5 genomes in RefSeq were considered. **B.** A phylogenetic tree linking previously sequenced *B. fragilis* reference genomes with sequenced colonies from two individuals (in red and blue). The effector genes encoded by each reference genome and the new sequenced genomes from stool are represented by colored squares as in Figure 1A.

**Figure S5.**
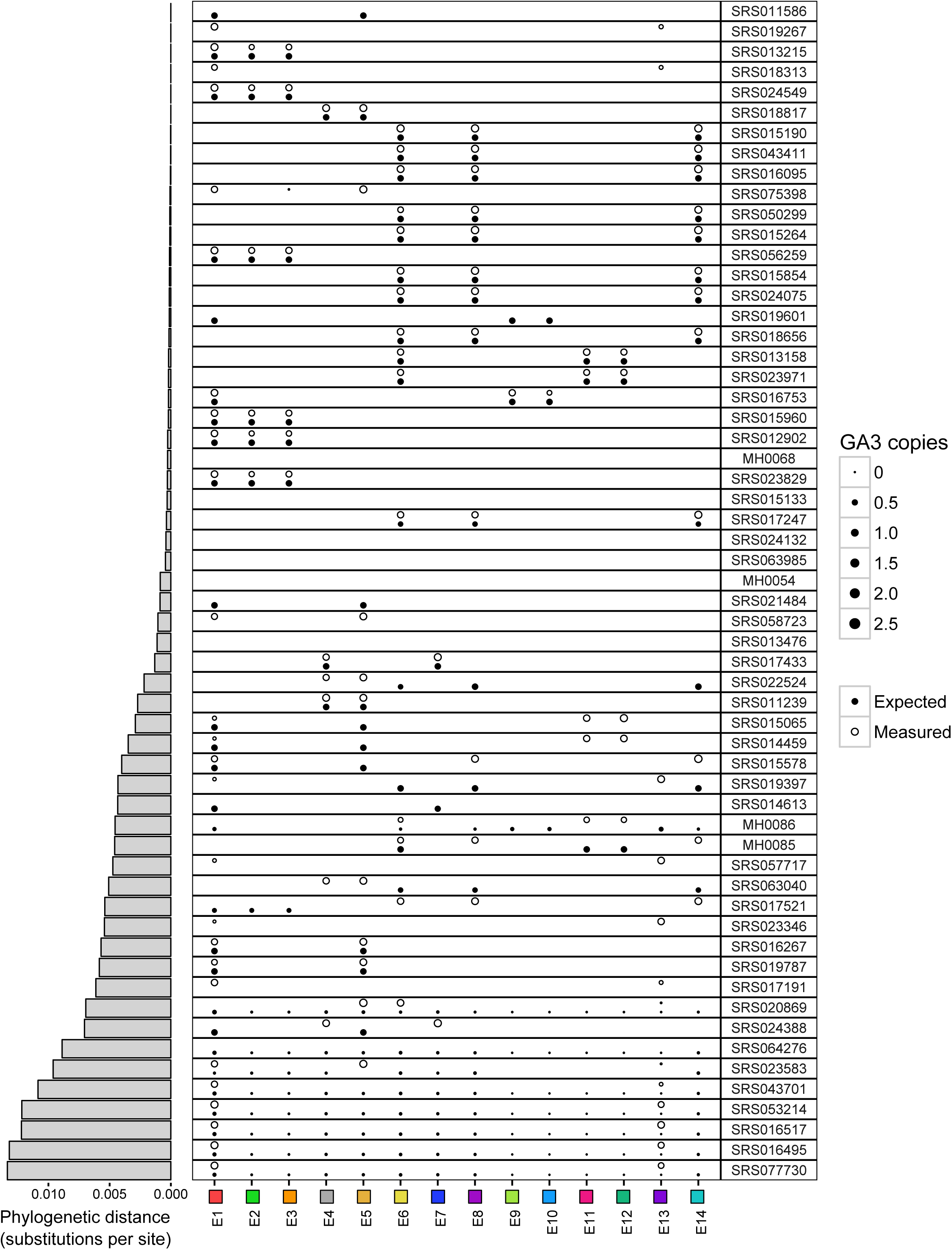
Measured and expected E–I gene abundances in HMP and MetaHIT samples. Measured abundances (hollow circles) are based on short read mapping to effector genes. Expected abundances (filled circles) are based on the reference strains phylogenetically closest to the inferred strain. Point size is scaled to the calculated copy number of each effector. The barplot on the left shows the phylogenetic distance between the inferred strain and its nearest reference strain in the phylogenetic tree.

**Figure S6.**
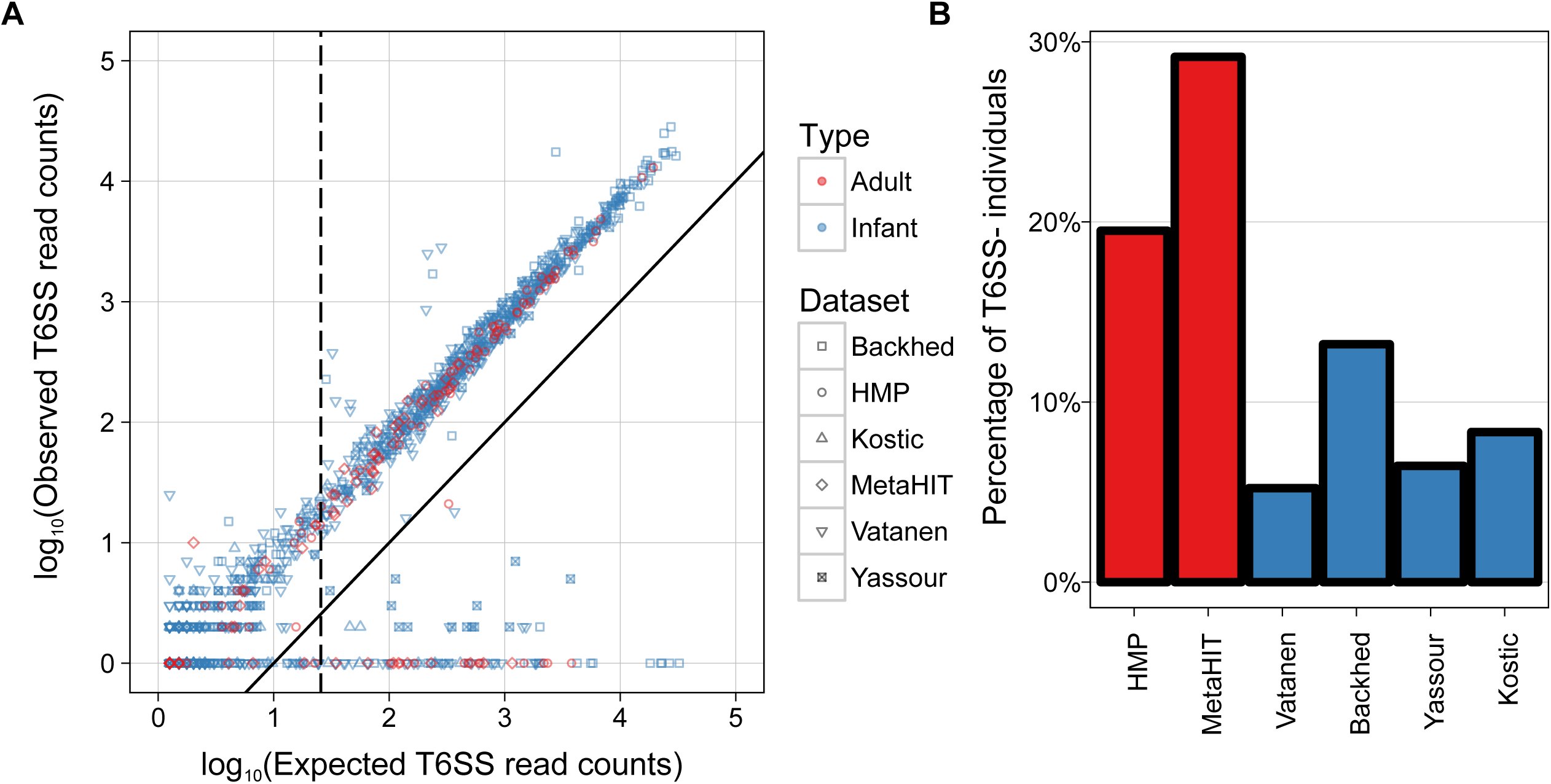
The prevalence of T6SS- samples in infants and adults. **A.** The relationship between the number of reads expected and measured to map to the GA3 T6SS structural genes, from each adult (red) and infant (blue) gut microbiome sample. The expected number is based on the number of reads mapping to *B. fragilis*-specific marker genes, normalized by gene lengths. The dotted line represents the cutoff used for determining that *B. fragilis* is present in a sample. The solid line represents an observed number of reads that is 10% of expected, and was used to distinguish T6SS+ from T6SS- samples. As evident by this plot, T6SS+ and T6SS- samples can be clearly defined. Different shapes correspond to different datasets, with adult samples colored in red and infant samples in blue. **B.** The percentage of individuals of those harboring *B. fragilis,* that lack the GA3 T6SS across different adult (red) and infant (blue) datasets. Individuals that were not consistent in terms of T6SS+/- classification across different time points were not included.

**Figure S7.**
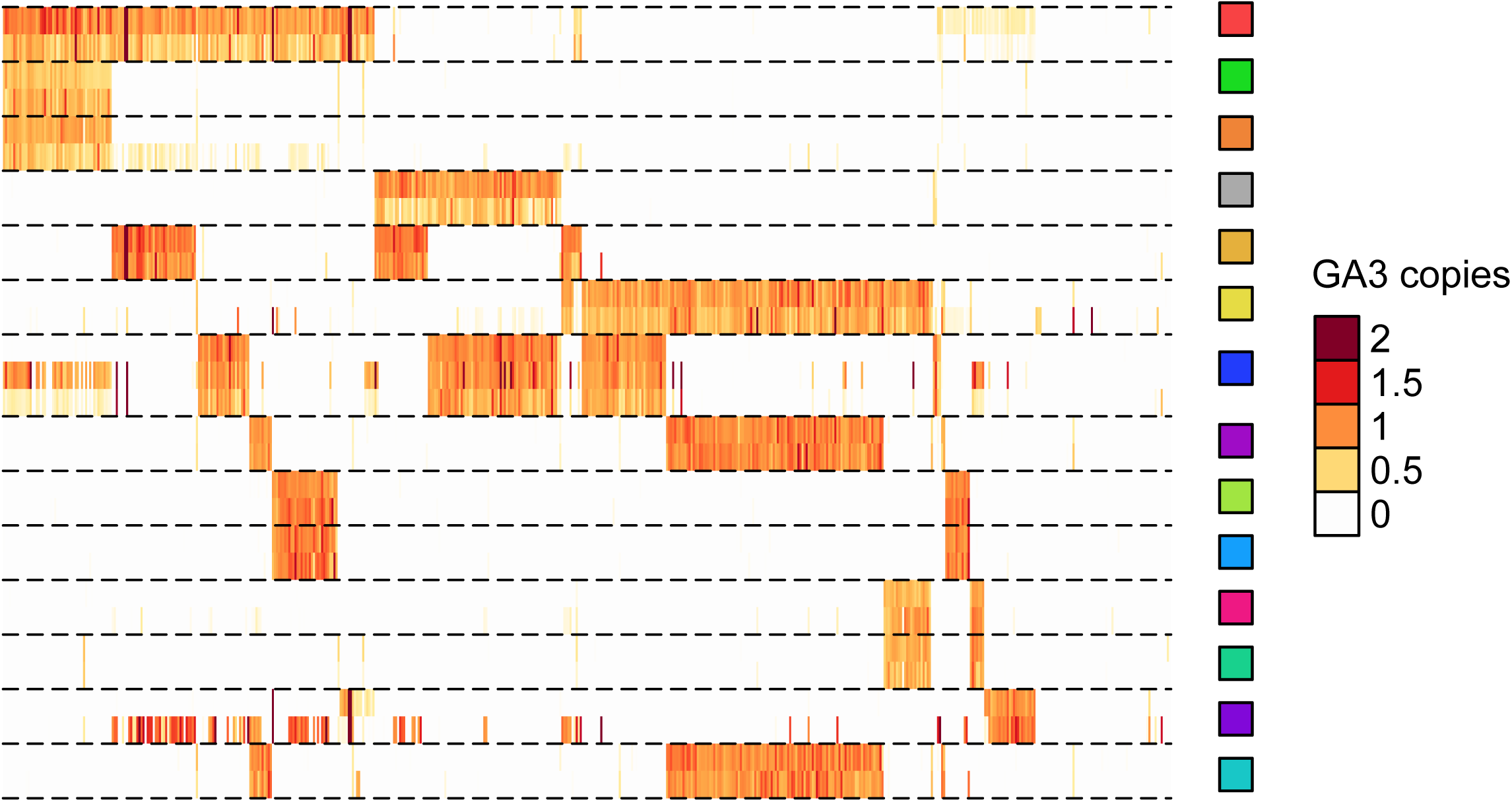
The abundance of *B. fragilis* GA3 E–I genes in infant microbiomes. Details and format are as in Figure 1D.

**Figure S8.**
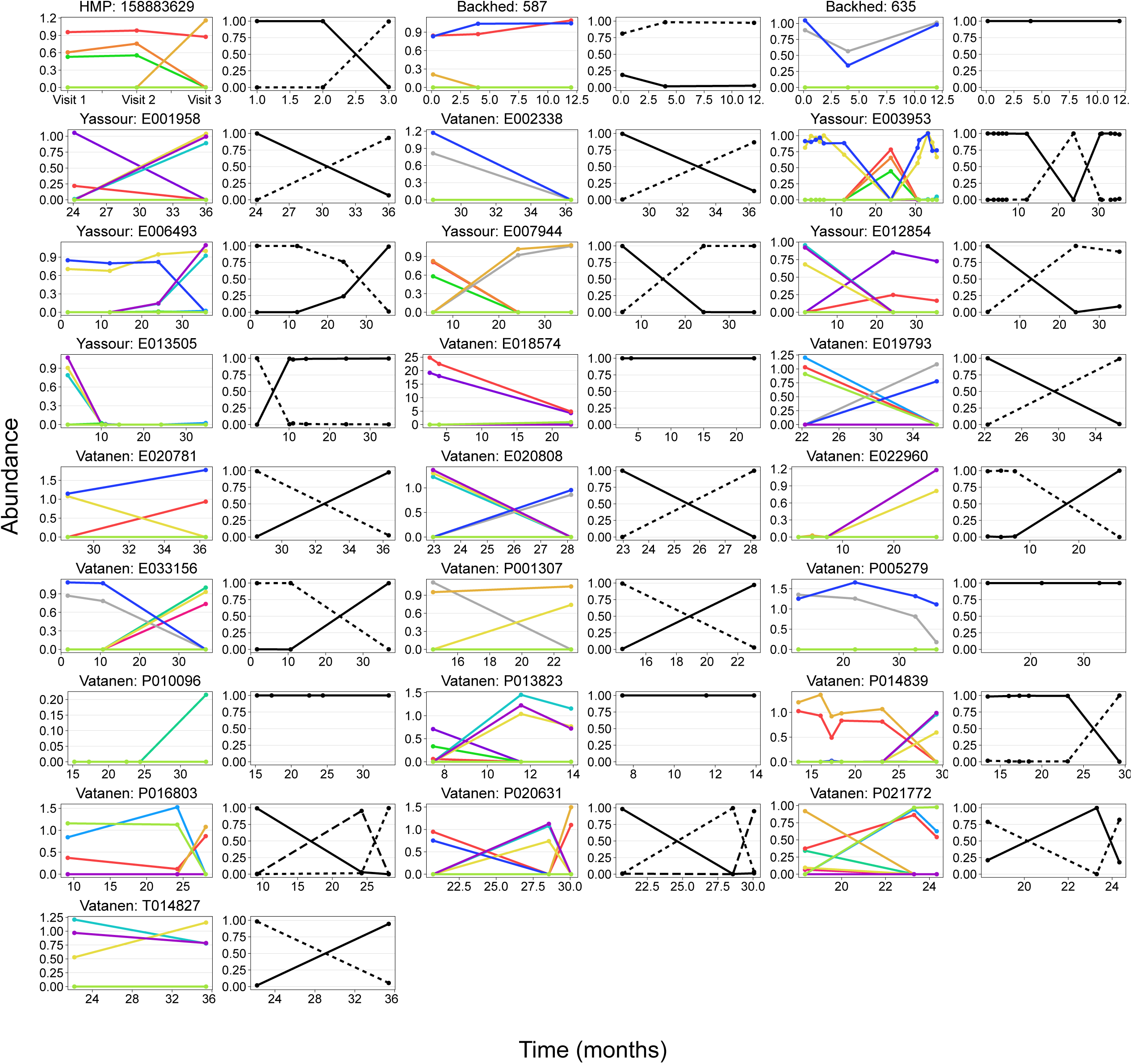
All E–I turnover and strain replacement events detected in adult and infant microbiomes. Details and format are as in Figure 3C-E (top plots). Each pair of plots is labeled with the individual code and the dataset it comes from.

**Table S1.** Genus-level differential abundance in T6SS+ vs. T6SS- samples.

| **Genus** | **Mean**<br>**Abundance**<br>**T6SS+ (%)** | **Mean**<br>**Abundance**<br>**T6SS- (%)** | **P value** | **FDR**<br>**Adjusted**<br>**P value** | **Passes**<br>**FDR** |
| --- | --- | --- | --- | --- | --- |
| *Faecalibacterium* | 3.8 | 10.6 | 8.3E-04 | 1.7E-02 | TRUE |
| *Bacteroides* | 57.9 | 37.5 | 3.1E-03 | 1.9E-02 | TRUE |
| *Ruminococcus* | 1.5 | 3.5 | 3.1E-03 | 1.9E-02 | TRUE |
| *Oscillospira* | 1.8 | 3.5 | 3.6E-03 | 1.9E-02 | TRUE |
| *Eubacterium* | 0.3 | 0.4 | 1.7E-02 | 7.2E-02 | FALSE |
| *Odoribacter* | 0.4 | 0.8 | 2.8E-02 | 9.8E-02 | FALSE |
| *Subdoligranulum* | 1.9 | 3.3 | 3.5E-02 | 1.1E-01 | FALSE |
| *Sutterella* | 2.4 | 0.1 | 7.6E-02 | 2.0E-01 | FALSE |
| *Dialister* | 1.2 | 0.7 | 1.1E-01 | 2.5E-01 | FALSE |
| *Clostridium* | 1.5 | 0.9 | 1.6E-01 | 3.2E-01 | FALSE |
| *Alistipes* | 5.7 | 6.7 | 1.8E-01 | 3.2E-01 | FALSE |
| *Coprococcus* | 0.5 | 0.6 | 1.8E-01 | 3.2E-01 | FALSE |
| *Akkermansia* | 0.1 | 0.3 | 2.2E-01 | 3.5E-01 | FALSE |
| *Lachnospira* | 0.7 | 0.7 | 2.6E-01 | 3.9E-01 | FALSE |
| *Parabacteroides* | 4.5 | 2.1 | 3.0E-01 | 4.2E-01 | FALSE |
| *Roseburia* | 1.5 | 2.7 | 4.5E-01 | 5.7E-01 | FALSE |
| *Blautia* | 0.9 | 0.5 | 4.6E-01 | 5.7E-01 | FALSE |
| *Megamonas* | 0.2 | 0.0 | 5.3E-01 | 6.2E-01 | FALSE |
| *Prevotella* | 0.1 | 0.3 | 6.0E-01 | 6.6E-01 | FALSE |
| *Phascolarctobacterium* | 0.8 | 0.8 | 8.0E-01 | 8.4E-01 | FALSE |
| *Escherichia* | 0.1 | 0.1 | 8.5E-01 | 8.5E-01 | FALSE |

